# Mapping the functional interactions at the tumor-immune checkpoint interface

**DOI:** 10.1101/2021.10.06.462889

**Authors:** Behnaz Bozorgui, Elisabeth K. Kong, Augustin Luna, Anil Korkut

**Affiliations:** Department of Bioinformatics and Computational Biology, UT MD Anderson Cancer Center, Houston, TX, 77030; Statistics, Rice University, Houston, TX, 77030, USA; Department of Data Sciences, Dana Farber Cancer Institute, Boston, MA 02215, USA

## Abstract

It is challenging to identify the tumor-immune system interactions that modulate immune states and immunotherapy responses due to the prohibitively complex space of possible interactions. Our statistical analysis framework, ImogiMap quantifies tumor-immune interactions based on their synergistic co-associations with immune-associated phenotypes. ImogiMap-based analyses recapitulated known interactions modulating immunotherapy responses and nominated the CD86/CD70 axis as an immunotherapy target that overlaps with *IFNG* overexpression and patient survival in endometrial carcinoma.

## Main

Genomically-targeted therapies and immunotherapies have led to improved patient survival in diverse cancer types (Waldman et al., 2020). Identification of biologically relevant tumor-immune interactions that mediate tumor cell survival may facilitate discovery of therapeutic targets particularly at the tumor-immune interface. The combination therapies which involve agents that target oncogenic processes and immune regulation, may induce more durable responses or even curative effects (Sharma and Allison, 2015). For example, expression of SERPINB9, a potential drug target and a member of the T-cell dysfunction signature genes, is upregulated in tumor cells by IFN*γ* in the tumor microenvironment (Jiang et al., 2018, McLaughlin et al., 2020). High expression of SERPINB9 confers resistance to CTLA4 checkpoint inhibition and therefore justifies the therapeutic benefit from co-targeting SERPINB9 and CTLA4. Despite such anecdotal findings, the interactions at the tumor-immune interface which inform effective therapies and immune states are relatively unexplored. It is indeed highly time and resource consuming to combinatorially search and identify tumor-immune interactions through cellular biology studies and even harder to determine which patient cohorts may benefit from an immunotherapy based on molecular signatures defined by such interactions. Existing tools such as TIMER and CIBERSORT have been very effective in deciphering the immune signatures in tumors as well as exploring interactions between immune features and expression of individual genes (Li et al, 2020, Chen et al., 2018). There are, however, no available software tools that investigate the combinatorial nature of interactions that potentially drive immune phenotypes at the tumor-immune interface.

Here, we introduce Immuno-oncology gene interaction Maps (ImogiMap), a method and informatics tool to automate the combinatorial searches for interactions between tumor-associated and immune checkpoint processes. Specifically, ImogiMap enables network analysis of combinatorial interactions between Immune checkpoint (ICP) genes, genes that are related to tumor-associated processes (TAP), and immune associated processes (IAPs). TAP genes may be any set of genes that are expressed in tumor ecosystems (likely but not necessarily in the tumor cells) and with functions involving hallmarks of cancer (e.g., proliferation, apoptosis, DNA repair, tumor metabolism, immune evasion) (Hanahan et al., 2011). IAPs may involve processes that are related to immunotherapy responses and immune states in tumor ecosystems (Blank et al., 2016) such as Immune cell infiltration (leukocyte fraction) (Hoadley et al. 2018), immune cell type fractions (Thorsson et al. 2018), and tumor mutation burden (Hoadley et al. 2018), together with mRNA-based pathway scores for epithelial mesenchymal transition (EMT) status (Mak et al., 2016), vascularization (Masiero et al., 2013), and IFNG expression (Gao et al., 2016). By mapping the ICP-TAP interactions, ImogiMap is able to inform on underlying mechanisms that jointly relate TAPs to therapeutically actionable ICPs and IAPs that are potential therapy response predictors. The outcome of the analysis is mechanistic links between tumors and the immune system that guide selection of possible prognostic markers and drug targets.

The minimal required input to ImogiMap is the mRNA expression data for ICP and TAP genes, and data (e.g., RNAseq, histological) that quantify IAP levels from matched samples. ImogiMap quantifies the individual interactions between pairs of TAP genes and ICP genes as synergy scores based on their co-association with the IAP of interest (Figure 1). To calculate synergy score, for each potentially-interacting TAP-ICP pair, the patient cohort is stratified into four sub-cohorts (low-low, high-low, low-high, high-high) based on TAP and ICP gene coexpression levels. The co-associations are quantified based on the IAP levels within the subcohorts. A non-additive deviation of the IAP levels within any one of the sub-cohorts with respect to other sub-cohorts indicates a synergistic interaction between the TAP and the ICP (see Methods and Figure 1). Note that ImogiMap does not rely on an a priori assumption involving a unidirectional relationship between TAP and ICP genes with the IAPs. A phenotype may be activated due to (or may be associated with) over-expression or loss of expression of a gene. As a result, the non-additive deviations in IAP levels may be observed in any of the four stratified sub-cohorts (see Figure 1 and methods) leading to a flexibility in analysis and interpretations of results.

**Figure 1.**
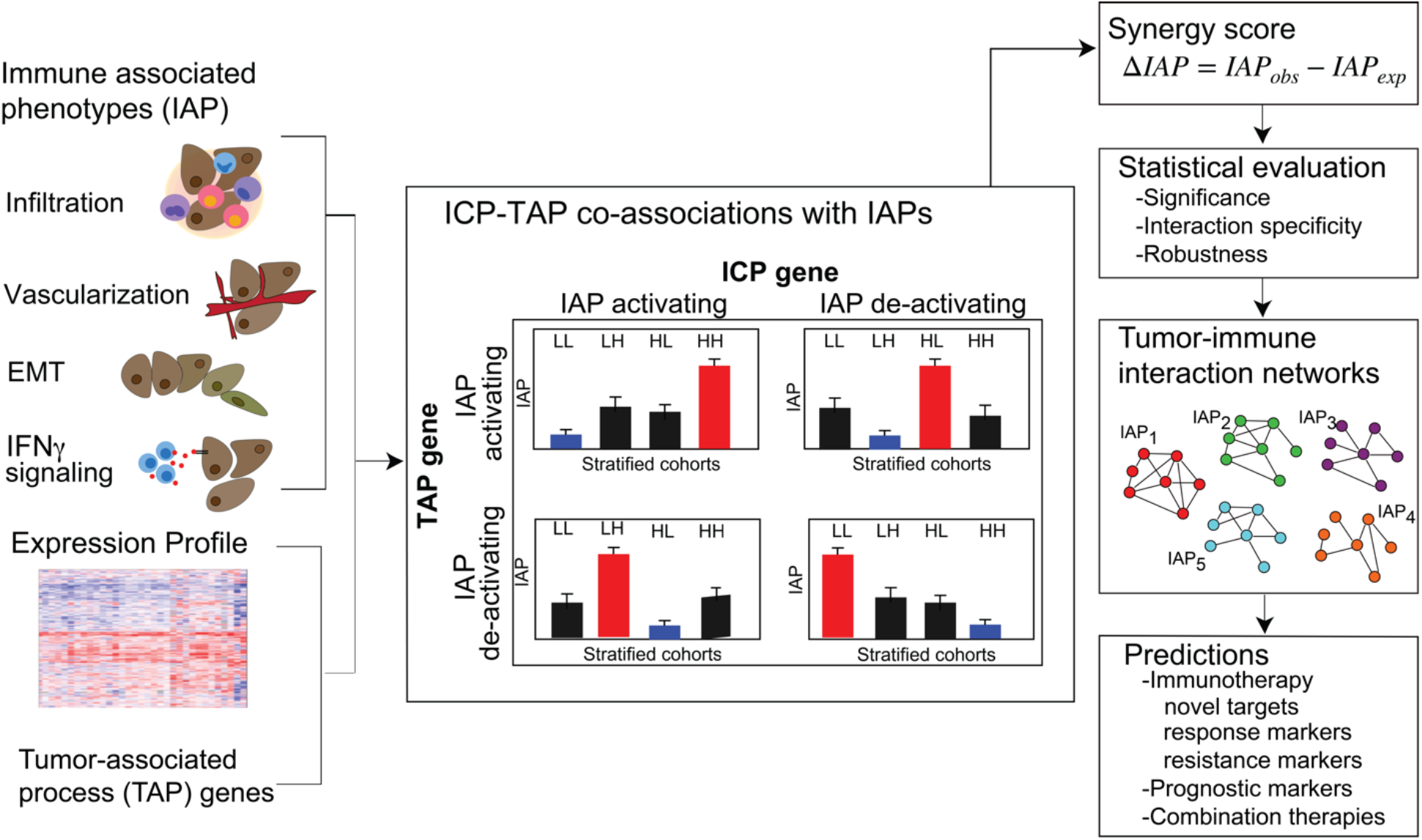
mRNA expression and immune-associated phenotype (IAP) levels are the inputs to ImogiMap. Based on the variation in IAP levels across patients, the algorithm calculates synergy scores between tumor-associated processes (TAP) that may constitute a functional gene signature and therapeutically actionable immune checkpoint (ICP) genes. For each TAP-ICP gene pair, the patient cohort is stratified into four sub-cohorts (LL, LH, HL, HH), and IAP levels are measured within each sub-cohort. A baseline sub-cohort (marked blue) and a target group (marked red) are determined based on four null assumptions on the relationship of the two genes with the IAP (both activating, both deactivating, and only one activating) (See methods). A synergy score under each assumption is calculated and their maximum is reported as the combinatorial association of the TAP-ICP pair with the IAP. For each IAP, a network of gene interactions from specific, robust, and significant interactions is constructed to reflect combinatorial relations between TAPs and ICPs in the context of an IAP.

To filter out potential false predictions; each synergistic interaction is statistically evaluated for robustness, statistical significance, and specificity. The robustness metric evaluates the stability of the interactions with respect to moderate changes in the exact data configuration. To assess the robustness, we compute the interaction synergies using data sets of partial cohorts that were randomly sampled from the complete cohort. The interaction robustness is quantified as the normalized root mean square deviation of synergy scores computed from the complete patient cohort vs. partial cohorts. The statistical significance is determined with a Wilcoxon signedrank test that compares the IAP measurements across the patient sub-cohorts based on the null hypothesis that the median IAP in a “target” sub-cohort and at least one of the remaining three sub-cohorts are sampled from an identical population (See figure 1 and methods for details). The interaction specificity is quantified by calculating the p-values of the synergy scores against a null distribution of scores for non-specific interactions. The null hypothesis for specificity evaluation is that the synergy score between a pair of TAP and ICP genes is equal to the synergy scores of TAP or ICP against other (randomly selected) genes. The null distribution of non-specific interactions is built as synergy scores between each of the genes of interest (TAP and ICP) and randomly sampled genes (N=1000) for a particular IAP. A robust synergy (not sensitive to moderate changes in data) with high significance (low p-value from a Wilcoxon signed-rank test) and high specificity (low p-value against a null-model of non-specific interactions) indicates a potentially functional interaction between tumor and immune genes for the IAP of interest.

Through integration of the significant, specific, and robust interactions, a collection of network models is constructed to map the associations of tumor-associated and potentially actionable immune processes with diverse immune phenotypes. Each network model maps multifaceted interactions between tumor and immune processes as well as ICP receptor-ligand interactions that lead to a significant change in a particular IAP. The collection of maps for diverse phenotypes presents a comprehensive atlas for comparative analysis among several IAPs. We also compare survival of patients whose tumors differentially express the interacting TAP and ICP genes to provide additional evidence on oncogenic relevance of phenotypic impacts. The resulting atlas informs on immune phenotypes, patient survival, and through interpretation of the maps, selection of potential immunotherapy options within patient cohorts carrying the tumor-immune interactions.

To validate our tool, we explored the interactions between T-cell dysfunction signature genes and ICPs at the tumor-immune interface. The T-cell dysfunction signature has been previously reported as a potential biomarker to predict response to immunotherapy in select cancer types including Uterine Corpus Endometrial Carcinoma (UCEC) (Jiang et al., 2018). Specifically, SERPINB9 which is part of the T-cell dysfunction signature stood up as a driver of immune checkpoint inhibitor resistance and is induced in response to IFN*γ* secretion. Here, we used a list of 26 core T-cell dysfunction signature genes in UCEC (Table S4), against 29 ICP genes in ImogiMap (Table S1). Significant, specific, and robust interactions that synergistically co-associate with *IFNG* gene expression are identified (Figure 2A–2C). Our analysis recapitulated the SERPINB9 as one of the top five T-cell dysfunction signature genes carrying strong interactions with ICP genes and associations with nonlinear increases in *IFNG* expression, as previously reported (Figure 2D). Together with SERPINB9, we find XCL1, a chemokine secreted by activated CD8 T-cells and NK cells, and CD5, a T-Cell receptor established as regulator of TCR and BCR signaling (Azzam et al., 2001), to be most frequently associated with *IFNG* levels through synergistic interactions with ICP genes.

**Figure 2.**
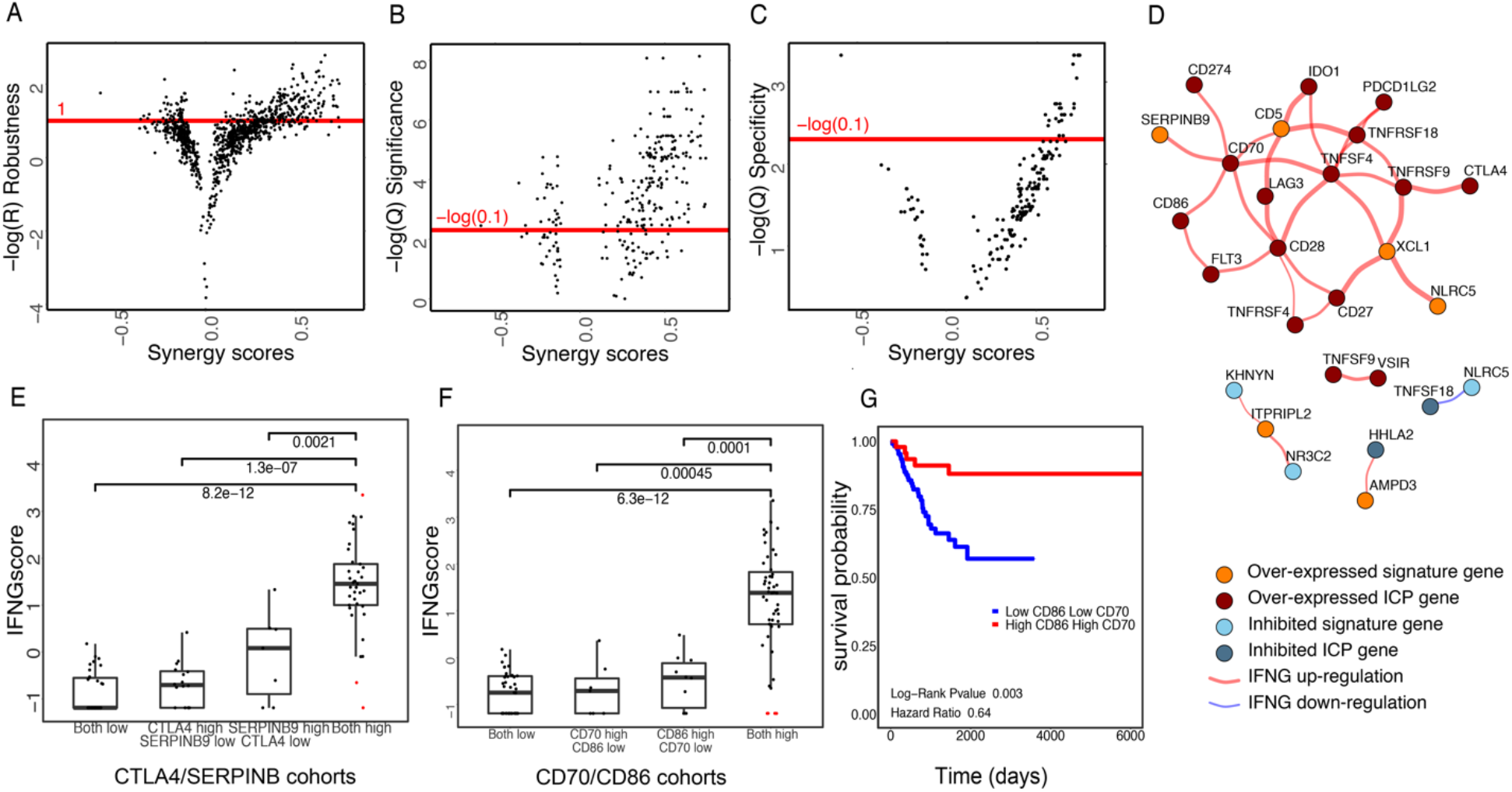
ImogiMap-based assessment of combinatorial interactions for T-cell dysfunction signature genes in Uterine Corpus Endometrial Carcinoma (UCEC), and therapeutically actionable ICPs, based on their associations with the IFNG expression. A) Synergy score robustness. A normalized root mean square deviation (RMSD) is calculated for each synergy score through random sampling (1000x) of sub-cohorts with 70% coverage of the complete cohort. Scores with high robustness, − *log(R)* > 1 (red horizontal line) are selected for further analysis (See methods). B) Synergy score significance. An FDR (Benjamini-Hochberg (BH) method) corrected *Q*-value based on Wilcoxon-signed rank test is calculated for robust scores. Scores with *Q*__*Significance*_ < 0.1 are selected for further analysis. C) A BH corrected Q-value for specificity is calculated for each robust and significant interaction (see methods). Scores with *Q*__*Specificity*_ < 0.1 are selected for further analysis. D) The graphical network representing robust, significant and specific combinatorial associations with IFNG levels. Red (Blue) edges represent synergistic upregulation (downregulation) of IFNG level. Dark red (Dark blue) vertices identify overexpression (inhibition) of ICP genes and orange (blue) vertices identify overexpression (low-expression) of T- cell dysfunction signature genes. E) Levels of IFNG expression in TCGA UCEC samples, stratified based on SERPINB9 and CTLA4 levels. P-values from Wilcoxon signed-rank test indicates statistical significance of differences in IFNG levels within the stratified sub-cohorts. G) Levels of IFNG expression in TCGA UCEC samples, stratified based on CD86 and CD70 levels. H) Kaplan-Meier survival curve for UCEC patients stratified by low/high expression of CD70 and CD86.

Encouraged by these results, we explored other interactions within the network models of tumor-immune interactions in UCEC (Figure 2D). We evaluated the association of each interacting gene pair in the networks with patient survival as assessed by p-values from a log-rank statistical test of survival differences. Among the 23 direct synergistic interactions, the CD70 and CD86 pair stood out as the factor that was most associated with progression free survival in UCEC patients (Figure 2F). CD86 and CD70 are immune co-stimulatory proteins that are expressed on antigen-presenting cells. Through interactions with their receptors (CTLA4, CD28, and CD27), both molecules promote T-cell activation and immune responses (Qin S. et al., 2019, Van de ven K. and Borst J., 2015, Hornero R.A. et al., 2020). The specific (*pvalue* < 0.05), robust (*R* < 0.05), significant (*pvalue* < 0.05) and high ranked synergy scores imply that expression levels of CD86 and CD70 synergistically co-associate with increased levels of IFN**γ** in UCEC patients (Figure 2G) supporting a role in immune stimulation. These findings posit the combinatorial targeting of CD86 and CD70 with agonist agents as a new candidate for immunotherapy with likely improved responses in UCEC patients with high CD86 and CD70 expression.

These findings demonstrate that ImogiMap generates novel hypotheses on (and confirms previously reported) relations between tumor associated processes and immune regulation at a scale not accessible easily by experimental methods. ImogiMap may help basic and translational researchers to discover novel immune-tumor interactions and potential vulnerabilities to combination therapies that target the immune-tumor interactions, improve the repertoire of actionable ICP targets, and identify patient cohorts that may respond to combination therapies based on their molecular signatures.

## Methods

### Immune checkpoints (ICPs) and Immune-associated phenotypes (IAPs)

Genes for 14 actionable ICPs as well as 15 additional genes of corresponding ligands or receptors are included (Table S1) based on a literature search and recommendations by the CRI clinical accelerator team (Qin S. et al., 2019, He X. et al., 2020, Yu J.X. et al., 2019, Marshall H.T. et al., 2018, Rotte A. et al., 2018, Cherif B., et al., 2021). Users have the option to input their own curated list of ICPs based on custom mRNA expression data or quantitative histologic assessments (e.g., IHC staining and quantification with relevant markers). In the absence of user-provided data, ImogiMap uses API functions in the curatedTCGAData package (Bioconductor), to access TCGA mRNA data. ImogiMap includes a list of 28 immune-associated phenotypes (Table S2). Levels of vascularization, EMT, and IFNG are each calculated using their corresponding signature gene lists (Masiero et al., 2013, Mak et al., 2016, Gao et al., 2016). For each of these phenotypes a pathway score is defined as the mean value of z-scaled mRNA expression levels of their corresponding signature genes in log scale. For TCGA-based analysis, precalculated values of immune cell infiltration and fractions of immune cell types (Table S3) are implemented based on DNA methylation (Hoadley et al. 2018) and CIBERSORT (Thorsson et al. 2018) assessment, respectively. Quantification methods, sources, and signature genes for immune phenotypes are listed in Table S2. Users have the option to include their own IAP values based on custom (omics or histology) data or TCGA data.

### IAP normalization and scaling

The measurements of IAPs are rescaled and normalized using a logistic sigmoidal function. The scaling facilitates comparisons of phenotypes and resulting synergy scores across different phenotypes. The rescaling is formulated to transform IAPs to a range of [0,1]. Three separate rescaling functions are formulated based on the initial range of IAP values across samples:

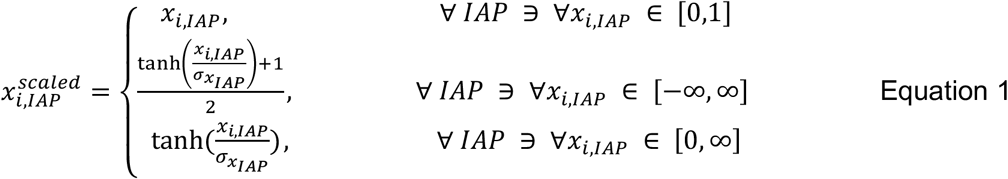

Where i is the sample index, IAP denotes a particular immune associated phenotype, X_i,IAP_ is the readout of the phenotype for sample i, σ_X,IAP_ is the standard deviation of readouts for IAP over all samples. The resulting X_i,AIP_^scaled^ has a dynamic range of [0,1] for any IAP and therefore cross-comparisons of metrics (e.g., synergy scores) calculated from scaled IAP values are possible. To demonstrate how the transformations affect phenotypes, the distribution of phenotypes before and after normalization are depicted in Figure S1.

### Calculation of gene pair synergies on IAPs

The synergy scores between immune checkpoint and tumor associated genes are calculated based on the non-linear deviation of the observed IAPs from the expected values for the condition that two genes are independently associated with the IAP. To measure synergy scores that quantify tumor-immune interactions, samples are first stratified into four groups based on median expression levels of two genes, as explained in the main text (i.e., “LL”, “LH”, “HL”, “HH”). First, to calculate a synergy score between two genes, a “baseline” sub-cohort and a target sub-cohort, which carry the presumably least and most co-associated IAP level respectively, are selected among the LL, LH, HL, or HH sub-cohorts. In the absence of a priori assumption on a relationship between expression of single genes and IAP levels (i.e., IAP-activating vs. IAP-deactivating genes), we analyze all possible configurations such that either the two genes may be activating/deactivating or have opposing relations with the IAP. For each configuration, the relevant baseline and target sub-cohorts with contrasting gene expression levels are identified and a synergy score is calculated as explained below. The configuration which results in the highest synergy score is selected as the metric for the potential combinatorial impact of the two genes on the IAP and is further analyzed.

The synergy score is quantified as follows: First, for each sub-cohort, the median deviation of IAP, *M*_*n*_, from the baseline is quantified as,

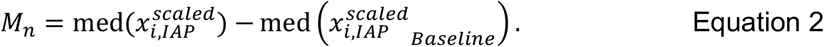

To reduce noise and ensure that the sign of *M*_*n*_ is correctly estimated, *M* values that are smaller than their corresponding standard error of the median, sem(*M*), are set to zero:

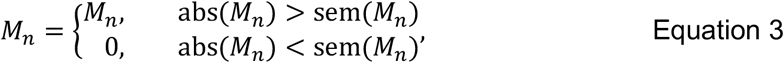

in which the standard errors are calculated using standard errors within each sub-cohort as,

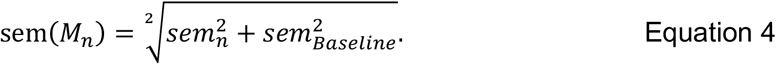

*M*_*E*_, an expected value for M_N_ in the target group, is calculated under the condition that each gene acts independently on the IAP. This is achieved through use of either of the two reference additivity models that are implemented in ImogiMap, Bliss independence model (Bliss C.I., 1939) or Highest Single Agent model (Berenbaum M.C., 1989). We define a synergy score, *S*, as the difference between the measured and the estimated *M* value in the target sub-cohort:

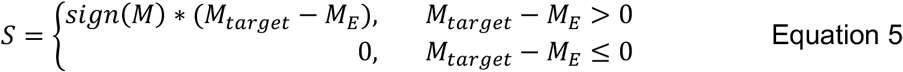

in which *sign(M)* is the direction of the change from the baseline in all sub-cohorts. Note that in the reference additivity models *M*_*E*_ for the target group can only be estimated if all three *M*_*n*_ values have the same sign, which indicates the same direction of change from baseline. If *sign(M)* differs in different sub-cohorts, no synergy score will be calculated, and a missing value will be reported. Finally, the synergistic interactions (*M*_*target*_ – *M*_*E*_ > 0) are selected for further statistical evaluation.

### Statistical evaluation of tumor-immune interactions

Through statistical assessment of the synergy scores between tumor-associated and immunoregulatory genes, we filter out the scores that are not robust, statistically significant, or specific.

**Robustness of synergistic interactions** to the underlying data configuration is tested through bootstrapping (1000x) of samples with partial coverage (default is 70% of a complete cohort), restratifying patient sub-cohorts based on the bootstrapped data, and calculating the synergy scores. The robustness score is defined as the normalized root mean square deviation of synergy scores between complete, *S*_*complete*_, and partial, *S*_*partial*_, datasets

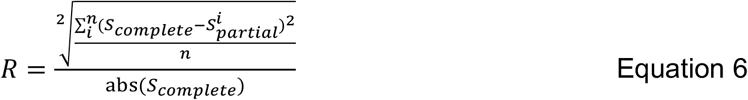

Where *S* is the synergy score, and *n* is the number of sampling to determine the *S*_*partial*_ set. Normalization with respect to absolute value of *S*_*complete*_ serves as the scaling factor to enable comparison of R value across diverse synergistic interactions. The least sensitive to the exact data configuration, therefore more robust and reliable scores, have lower *R* and are assessed by ranking the sensitivity scores for all interactions.

**The statistical significance of interactions** is assessed with a Wilcoxon signed-rank sum test comparing the median values of immune phenotypes in the target sub-cohort and the remaining three sub-cohorts (LL, LH, HL, HH). In the statistical assessment, the null model is that the statistical median values of immune phenotypes in the target sub-cohort and at least one of the remaining three sub-cohorts are equal. The maximum of the three Wilcoxon p-values is reported as the p-value of the interaction. The subcohort IAP values are compared on boxplots which contains Wilcoxon p-values for group comparisons.

**The specificity of a synergistic interaction** between a TAP and ICP gene pair is quantified. The specificity implies that the observed synergy score between the two genes of interest is significantly higher than the synergy scores of each of the gene pair against other genes in the genome. First, two separate null models are generated by calculating the synergy scores of each of the two genes against a set of genes that are randomly sampled from the genome (default *N*_*genes*_ = 1000). For each of the two genes, a P-value is calculated against the two null models. The highest of the two P-values, *P*_*max*_, which indicates the lowest specificity is used to assess the specificity of the interaction against the whole genome. A low P-value (*P*_*max*_ < 0.05) indicates high specificity.

### Network models of synergistic immune-tumor interactions

The synergistic interactions that are significantly strong, robust and specific are selected to construct a network model which captures the combinatorial interactions between the tumor-associated and immune genes. In the network models, the edges represent interactions between gene pairs for which a significant, specific and robust synergy on IAP exists. Weak interactions are also filtered out based on a user-defined cut-off value. The network model is visualized using the igraph application (Csardi and Nepusz, 2006).

## Availability

ImogiMap R package is available at https://github.com/korkutlab/Imogene

## Acknowledgement

AK is supported with grants from MDACC Support Grant P30 CA016672 (the Bioinformatics Shared Resource), OCRA Collaborative Research Award, CPRIT High-Impact/High-Risk Award (RP170640), NIH/NCI U01CA253472-01A1.

## COI

The authors declare no competing financial interests.

## Contributions

A.K. and B.B. designed and executed the project. A.K., B.B., E.K.K. and A.L. analyzed the results and wrote the paper. B.B. developed the software package.

## Supplementary information

**Table S1.** Immune checkpoints from a literature search and Cancer Research Clinical Accelerator, and molecules that interact with them as ligands or receptors.

| Gene name | Protein name | Potential interactions |
| --- | --- | --- |
| PDCD1 | PD-1 | PD-L1 , PDL2 |
| CD274 | PD-L1 | PD-1 |
| CTLA4 | CTLA-4 | CD28, B7-1, B7-2 |
| CSF1R | CSF1R | CSF1 |
| TNFRSF9 | 4-1BB | 4-1BBL |
| TNFRSF18 | GITR | GITRL |
| CD27 | CD27 | CD70 |
| ICOS | ICOS | B7-h2 |
| TNFRSF4 | OX40 | OX40LG |
| PDCD1LG2 | PD-L2 | PD-1 |
| CD70 | CD70 | CD27 |
| CSF1 | CSF1 | CSF1R |
| TNFSF9 | 4-1BBL | 4-1BB |
| TNFSF4 | OX40LG | OX40 |
| TNFSF18 | GITRL | GITR |
| CD28 | CD28 | CTLA-4 |
| CD80 | B7-1 | CTLA-4 |
| CD86 | B7-2 | CTLA-4 |
| ICOSLG | B7-h2 | ICOS |
| CD276 | B7-h3 |  |
| VTCN1 | B7-h4 |  |
| VSIR | B7-h5 |  |
| NCR3LG1 | B7-h6 |  |
| HHLA2 | B7-h7 |  |
| LAG3 | LAG-3 |  |
| IDO1 | IDO1 |  |
| FLT3 | FLT3 |  |
| NT5E | CD73 |  |
| TLR3 | TLR-3 |  |

**Table S2.** List of Imogene’s default immune-associated phenotypes

| Phenotype | Source |
| --- | --- |
| EMT score | mRNA expression of EMT signature genes (Mak et al., 2016) |
| Vascularization | mRNA expression of Angiogenesis signature genes (Masiero et al., 2013) |
| IFNG score | mRNA expression of IFNG (Gao et al., 2016) |
| Non-silent tumor mutation burden per MB | Whole exome or genome sequencing (Hoadley et al. 2018) |
| Silent tumor mutation burden per MB | Whole exome or genome sequencing (Hoadley et al. 2018) |
| Leukocyte fraction | DNA methylation (Hoadley et al. 2018) |
| 22 immune cell type fractions | CIBERSORT analysis of mRNA expression (Thorsson et al. 2018) |

**Table S3.** List of 22 immune cell types from CIBERSORT

|  |
| --- |
| Memory B Cells |
| Naive B Cells |
| Plasma Cells |
| CD8 T cells |
| Naive CD4 T cells |
| Activated memory CD4 T cells |
| Resting Memory CD4 T cells |
| Follicular Helper T Cells |
| Regulatory T Cells |
| Gamma Delta T Cells |
| Activated Natural killer Cells |
| Resting Natural Killer Cells |
| Monocytes |
| M0 Macrophages |
| M1 Macrophages |
| M2 Macrophages |
| Resting Dendritic Cells |
| Activated Dendritic Cells |
| Activated Mast Cells |
| Resting Mast Cells |
| Eosinophils |
| Neutrophils |

**Table S4.**
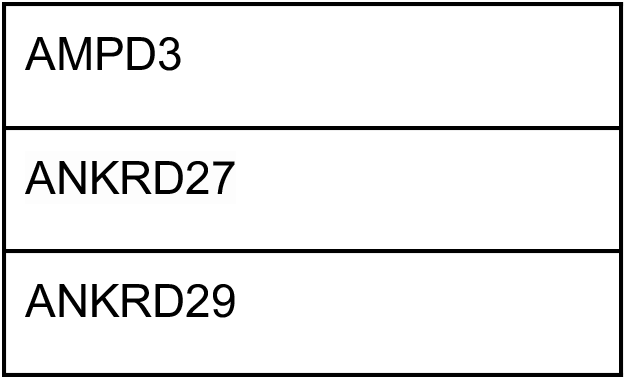
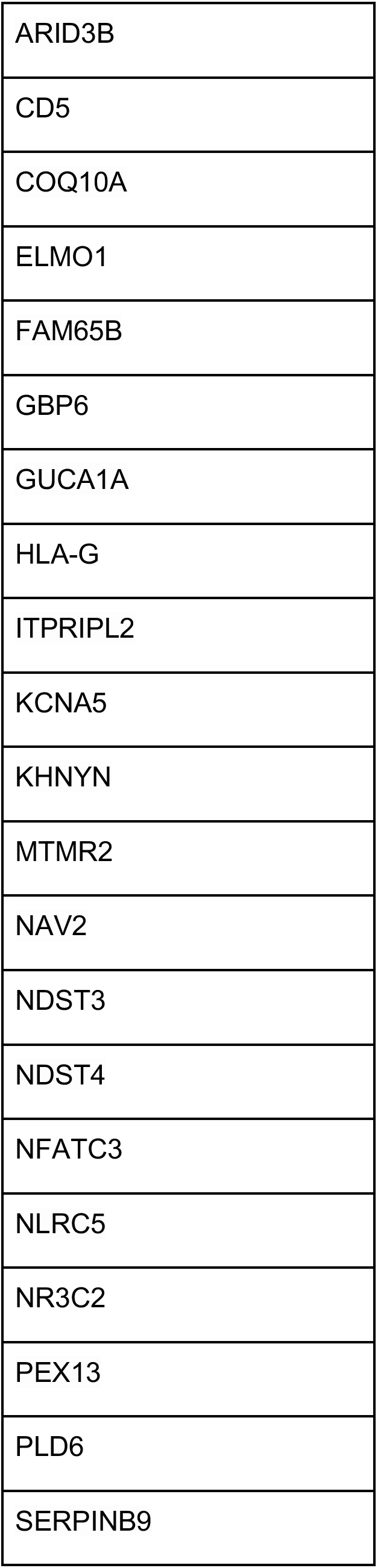
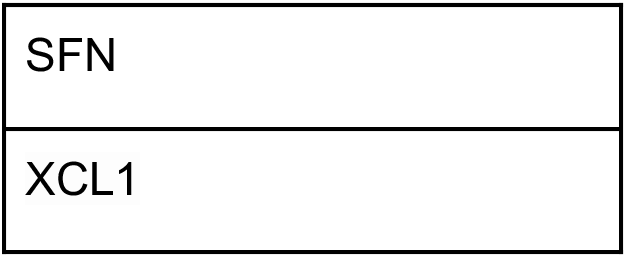
List of T cell dysfunction signature genes (Jiang et al., 2018)

**Figure S1.**
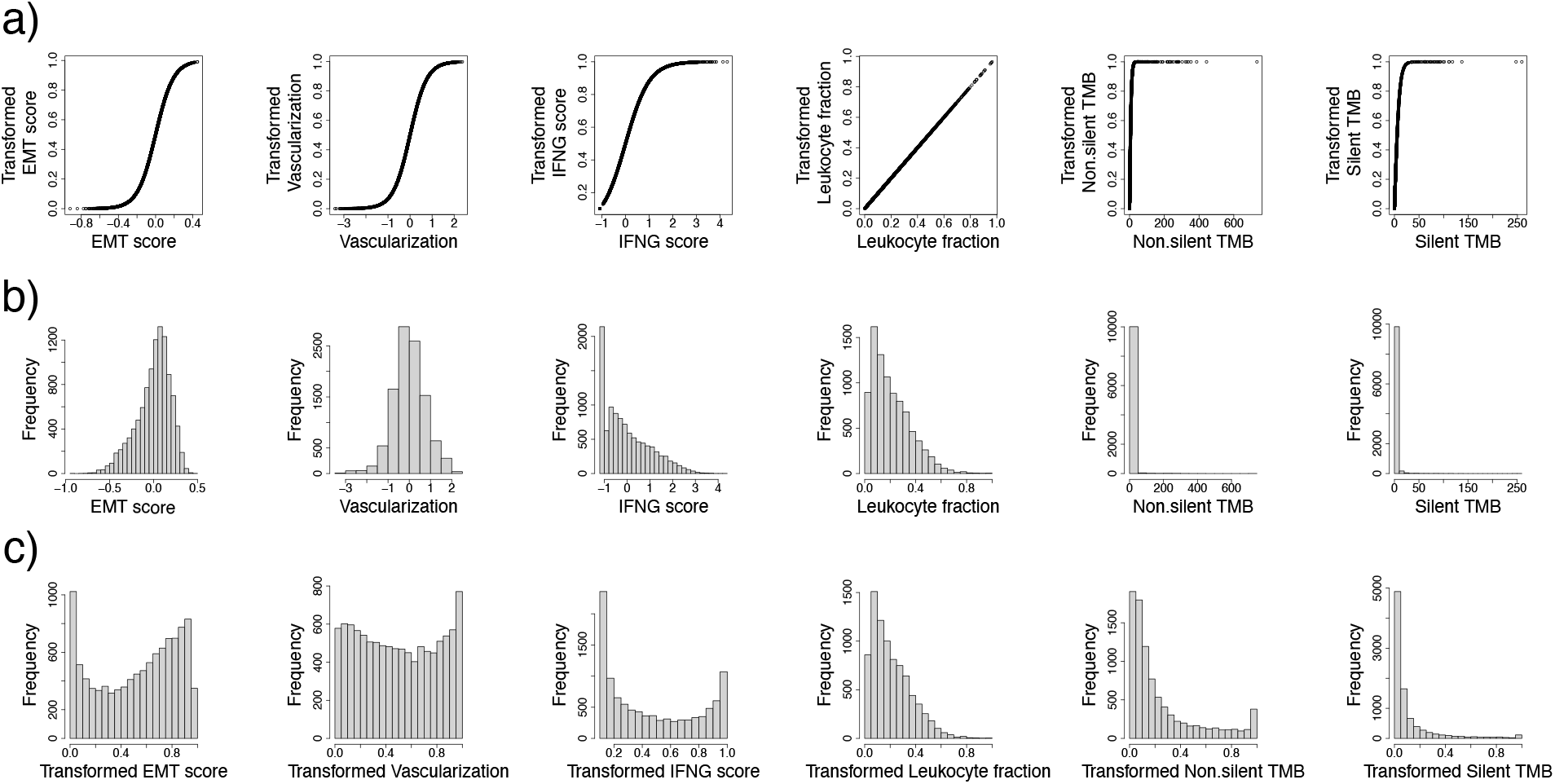
a) Graphs of IAP transformation functions. b) Distribution histograms of IAP values in PanCan TCGA data. c) Corresponding distribution histogram of transformed IAP. Note that Leukocyte_fraction and fractions of immune cell types do not change under transformation, as they are inherently positive and smaller than 1.

## Notes

### Competing Interest Statement

The authors have declared no competing interest.

